# Liver regeneration by oval cells employing bistability of stemness-senescence, Hippo signaling, EMT-MET, and polyploidy circuit

**DOI:** 10.1101/2024.03.26.586724

**Authors:** Marija Lazovska, Kristine Salmina, Dace Pjanova, Bogdan I. Gerashchenko, Jekaterina Erenpreisa

**Affiliations:** Institute of Oncology and Molecular Genetics, Riga Stradins University, Riga, Latvia; Riga Stradins University, Riga, Latvia; Latvian Biomedical Research and Study Centre, Riga, Latvia; R.E. Kavetsky Institute of Experimental Pathology, Oncology and Radiobiology, National Academy of Sciences of Ukraine, Kyiv, Ukraine

**Keywords:** Liver regeneration, oval cells, bipotential progenitor, stemness-senescence, EMT-MET, polyploidy circuit, turbulence, replication stress, Hippo pathway

## Abstract

Liver hepatocytes possess remarkable regenerative capabilities, yet severe damage may compromise this process. Liver progenitor (“oval”) cells exhibit the potential to differentiate into both hepatocytes and cholangiocytes, making them promising candidates for cell therapy. However, their mechanisms in liver regeneration are not clear. Here, on rat liver oval stem-like epithelial cells (WB-F344) a wound healing assay was performed. The scratched near-confluent monolayers (70% area removed) underwent the G1-arrest, bi-nucleation at 10-12 hours post-wounding, starting movement of epithelial to mesenchymal transition (EMT) cell portion into the wounded areas. Nanog nuclear upregulation, fragmentation, and transition as granules into cytoplasm and around, along with p16^Ink4a^ nuclear intrusion from the cytoplasm, loss of epithelial markers, and YAP1/Hippo activation were seen near the wound edge. The replicative stress and proliferation boost followed, documented at 24 hours. Proliferation concluded at 40-48 hours, accomplished by reconstitution of epithelial tissue, the disappearance of Nanog granulation and p16^Ink4a^ return to the cytoplasm, releasing excess. This investigation reveals novel regulatory facets in liver regeneration by oval cells. It accentuates the stemness-senescence bistable switch regulated by reciprocal nucleo-cytoplasmic transitions of opposite regulators, coordinated with Hippo-pathway switch, replicative stress, and boost, along with ploidy, EMT-MET and paracrine secretome circuits - enabling successfully resolving the massive injury.

Fig 1.
Graphical abstract.
Bistable nuclear-cytoplasmic switch between stemness and senescence regulators in the wound healing process by oval liver cells: (1-2) Priming phase: (1) at the wound edge, (2) in the wound; (3) Proliferative phase, wound closure. Nanog – green; p16INK4A – red, EMT cell - with blue nucleus.

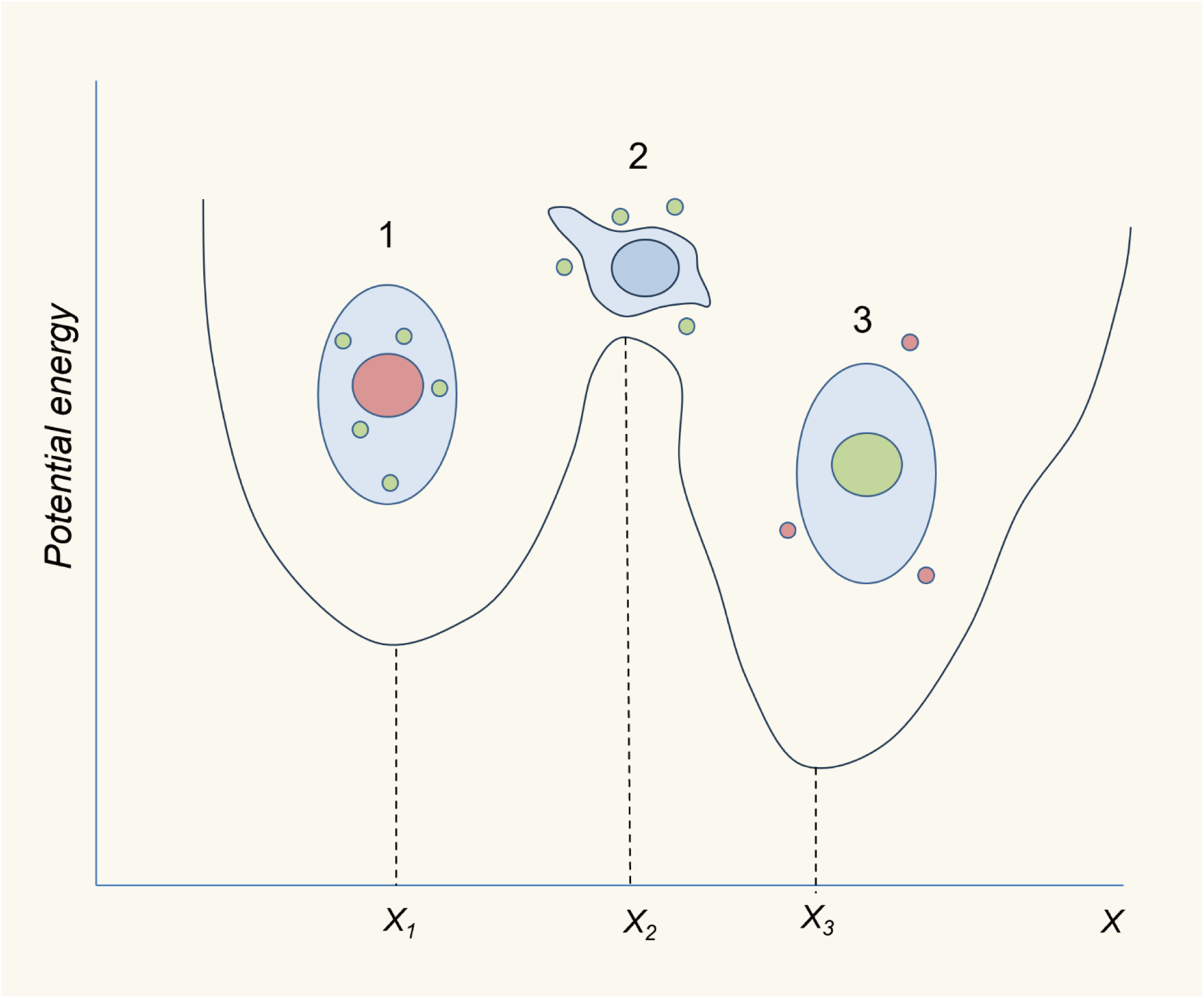

## Introduction

The mammalian liver possesses a remarkable capacity to regenerate after the injury while recovering its volume and functionality. However, the regenerative capacity of the liver is not endless: while the removal of up to two-thirds of the liver (partial hepatectomy; PH) introduced in the experiment by Higgins and Anderson in 1931 [1] can be safely repaired, larger resection is at risk of liver failure and following the death of the patient. Unfortunately, there could be life-threatening situations due to progressive complications of liver pathology that need to be eradicated as soon as possible by surgical means. According to statistics, hepatic disease is one of the top causes of death globally and there is an ascending tendency of diseases such as Hepatitis B and C often ending in cirrhosis and liver cancer [2]. At present, stem cell therapy is being extensively explored to tackle prognostically unfavorable liver diseases [3]. Despite the progress, the challenges remain, and yet liver transplantation remains the only curative treatment for patients with end-stage liver diseases.

Liver progenitor cells (LPCs)-driven liver regeneration was shown to be activated in case of massive liver injuries (Duncan et al., 2009). It involves the activation and proliferation of the LPCs expressing developmental markers and having the potential to become both mature hepatocytes and biliary cells [4]. The events induced after the typical liver injury are usually categorized into three phases: priming (activation), proliferation and termination [5]. Different signaling pathways and molecular processes are involved, both LPCs- and hepatocyte-driven, also called typical liver regeneration performed by a healthy liver. Many pathways are shared by both types of regeneration, but LPCs-driven repair remains less understood. The Hippo pathway, one of the central growth regulators, was shown to accompany the whole healing process from cell activation to the termination of response [6]. This pathway is involved in replicative stress [7], and cellular polyploidy, which is described in normal liver and boosted by regeneration [8, 9]. Senescence is another mechanism involved in liver repair; however, its role is controversial and poorly understood [10, 11]. The unique hepatocyte plasticity and reprogramming to an early postnatal-like state in response to injury were shown using a single-cell RNA sequencing [12].

In this work, using an *in vitro* wound healing assay, we study rat LPCs (known as “oval” epithelial cells) in response to mechanical injury. The primary goal of the study is to find out how the previously described mechanisms are interconnected in time and space by combining phenotypical and molecular studies of LPCs throughout the healing process. Here, in addition to the already known “polyploidy circuit”, in which polyploid hepatocytes undergo ploidy reduction during injury response and subsequent re-polyploidization [13] together with Hippo pathway on-off state, we describe cooperation and “competition” between the opposite regulators of cell fate - senescence/quiescence and stemness/self-renewal. Replicative stress, which is likely imposed by YAP1 in this wound healing assay, could be a key contributor in this bistability directed towards the regulation of cell-fate changes.

## Methods

### The cell line

The non-transformed male diploid WB-F344 rat liver epithelial cell line [14] was kindly provided by Dr. Tomonori Hayashi (Radiation Effects Research Foundation, Hiroshima, Japan). It is a benign wild-type p53 stem-like cell line, rapidly proliferating, and potentially capable of differentiating into both hepatocytes and cholangiocytes. In addition, these cells are relatively radioresistant as they do not undergo rapid and massive cell death even after high-dose irradiation up to 10 Gy [15]. The cells were cultured in RPMI 1640 growth medium (Sigma-Aldrich, St. Louis, MO, USA) supplemented with 7% fetal bovine serum (FBS) (Sigma-Aldrich, St. Louis, MO, USA) and 600 μg/ml L-glutamine (Gibco; Thermo Fisher

Scientific, Waltham, MA, USA). Cells were incubated at 37°C in a humidified 5% CO2 atmosphere with complete medium replacement performed every 24 h. Cells were routinely subcultured after reaching 70-80% confluency every three days.

### Wound healing (scratch) assay

Before wound healing assay, cells were grown on 4-well glass chamber slides coated with poly-D-lysine (Thermo Fisher Scientific, Waltham, MA, USA). About 70,000 cells were seeded in each chamber well with a total aliquot of growth medium of 1 ml. To mimic the organ regeneration after the wounding and to test the wound-induced plasticity of WB-F344 cells, a simple wound healing (scratch) assay was applied. For this purpose, cell monolayers that were maintained in chamber slides reaching the near- confluence of approximately 95% after 24–30 h culturing were used. A sterile 1000 μl pipette tip was used to scratch a grid composed of longitudinal and transverse lines (approximately 9 and 6 lines, respectively), so the distance between the closest non-intersecting lines was almost equal (∼ 2 mm). Approximately two-thirds of the monolayer was removed. To take away cell debris, the cell monolayers were washed three times with pre-warmed culture media by gentle shaking followed by the addition of fresh media. Cells were then incubated at the aforementioned conditions until the desired time point or full closure of the wound. An Experimental design of the wound healing assay is shown in Fig 2.

**Fig 2.**
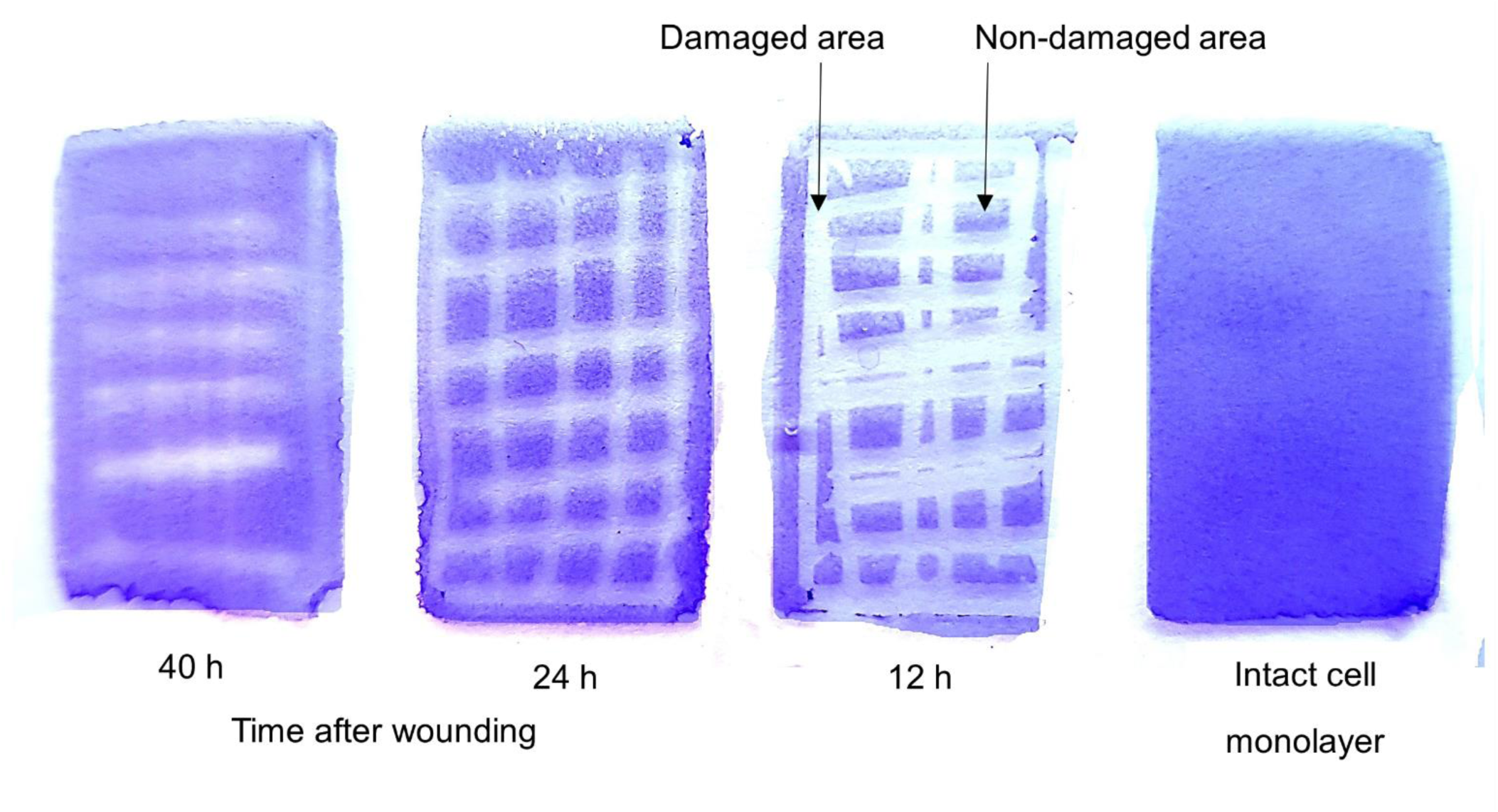
Experimental design. Monolayers of WB-F344 cells after reaching near 100% confluence in chamber slides were scratched and at 40, 24, and 12 h after wounding were fixed and stained with Toluidine blue (TB) for analysis. The highlighted enlarged region of interest from one of the TB-stained monolayers demonstrates different densities of cells in damaged (scratched) and non-damaged (non- scratched) areas colored in white and blue, respectively. Based on spatial localization to wound the edge the cells were subsequently analyzed for DNA content by image cytometry and cell-cycle phase- dependent nuclear morphology.

### Immunofluorescence staining

For immunofluorescence staining the chamber slides with live scratched adherent cells were removed from growth media and rinsed three times with warm phosphate-buffered saline (PBS), pH=7.4 (Sigma- Aldrich, St. Louis, MO, USA), and afterwards a droplet of warm FBS was added to each well. Before the next manipulations, a chamber was gently pulled off from the slide and the slides were allowed to shortly air-dry vertically. Then cells were fixed in methanol for 7 min at -200C, dipped 10 times in ice-cold acetone, and allowed to briefly dry. Slides were washed three times in 0.01% tris-buffered saline/Tween 20 (TBST) for 5 min each. Subsequently, they were blocked for 40 min in 1% bovine serum albumin (BSA) in PBS with 0.05% Tween-20 (Sigma-Aldrich, St. Louis, MO, USA) at room temperature (RT). After blocking, the samples were covered with 1% BSA in PBS with 0.025% Tween-20 containing primary antibodies and incubated overnight at 40°C in a humidified chamber. The next day, slides were washed five times in 0.01% TBST and covered with TBST containing appropriate secondary antibody in 1:300 dilution (goat anti-mouse IgG Alexa Fluor 488, goat anti-rabbit IgG Alexa Fluor 594 (Invitrogen, Carlsbad, CA, USA)) and incubated for 40 min at RT in the dark in a humidified chamber. After incubation, the samples were washed again five times with 0.01% TBST and once for 2 min in 1xPBS. Cells were counterstained with DAPI (0.25 μg/ml) for 2 min and embedded in Prolong Gold (Invitrogen, Carlsbad, CA, USA). When staining for α-tubulin and actin, the cells were fixed with fresh warm (∼ 37°C) 4% paraformaldehyde for 15 min at RT, then washed three times with 1xPBS and permeabilized with 0.5 % Triton-X100 (Sigma-Aldrich, St. Louis, MO, USA) in 1xPBS for another 5 min, washed again; and further the previously described steps starting from blocking were carried out. Three independent experiments for each combination of primary antibodies were performed. For microscopic observations and sample analysis, a fluorescent light microscope (Leitz Ergolux L03-10, Leica, Wetzlar, Germany) equipped with a color video camera (Sony DXC 390P, Sony, Tokyo, Japan) and RGB filter-set were used. Primary antibodies and their sources are listed in Table 1.

**Table 1.** List of antibodies used in this study for immunofluorescence staining.

| Antibody Against | Description | Used concentration | Product no. and Manufacturer |
| --- | --- | --- | --- |
| **α-Tubulin** | Mouse monoclonal | 1:200 | T5168, Sigma-Aldrich, St. Louis, MO, USA |
| **Albumin** | Rabbit polyclonal | 1:200 | PA5-85166, Invitrogen, Carlsbad, CA, USA |
| **Nanog** | Mouse monoclonal | 1:50 | N3038, Sigma-Aldrich, St. Louis, MO, USA |
| **Twist1** | Mouse monoclonal | 1:100 | sc-81417, Santa Cruz, Dallas, TX, USA |
| **Cytokeratin 7** | Mouse monoclonal | 1:100 | MA1-06315, Invitrogen, Carlsbad, CA, USA |
| **Oct 3/4** | Mouse monoclonal | 1:40 | sc-5279, Santa Cruz, Dallas, TX, USA |
| **p16<sup>Ink4a</sup>** | Rabbit polyclonal | 1:50 | ab84075, Abcam, Cambridge, UK |
| **TEAD1** | Mouse monoclonal | 1:200 | GT13112, Invitrogen, Carlsbad, CA, USA |
| **F-ACTIN** | Phalloidin-iFlour 594 Conjugate | 1:300 | ab176757, Abcam, Cambridge, UK |
| **YAP1** | Rabbit polyclonal | 1:500 | PA-46189, Invitrogen, Carlsbad, CA, USA |
| **c-Myc** | Mouse<br>monoclonal | 1:100 | sc-40, Santa<br>Cruz, Dallas, TX,<br>USA |

### DNA staining and image cytometry

To evaluate DNA amount in cells, count mitotic figures and estimate cell cycle dynamics the samples were stained with toluidine blue (TB) by the method [16], by the following protocol. Chamber slides with adherent scratched or control cells were washed three times with 1xPBS as described before, with subsequent addition of warm FBS and air-drying for 30 min. Then slides were fixed in ethanol/acetone (1:1) for at least 30 min at 40°C and then air-dried again. Next, slides were hydrolyzed with 5 N HCl for 20 min at 20-21°C. In some experiments, to partly preserve RNA staining and observe nuclei together with the cytoplasm, a shortened hydrolysis for 10 sec was performed. Slides were then washed in distilled water five times 1 min each and stained with 0.05% TB in 50% citrate-phosphate McIlvain buffer at pH=4 for 10 min. They were briefly rinsed in distilled water and blotted with filter paper. Subsequently, the slides were dehydrated by incubating twice in butanol for 3 min each at 37°C, cleared twice in xylene for 3 min each at RT and embedded in DPX. Ready samples were imaged in the green channel using a Sony DXC 390P color-calibrated video camera (Sony, Tokyo, Japan). From the greyscale images, DNA content was measured as the IOD using Image-Pro Plus 4.1 software (Media Cybernetics, Rockville, MD, USA). Cell DNA content or ploidy number was compared to IOD values for metaphases and anaphase- telophase (ratio= 2) and equated to diploid (2C) DNA values in G1 cells. A sum error of 10% was established for this DNA stoichiometry method using in situ image analysis, and this interval was considered for estimating proportional distribution of cell-cycle phases. For cell cycle measurements, approximately 300-500 interphase cells were collected at each experimental time point. Mitotic indexes were measured microscopically per 1000 cells, as well as other characteristics, such as the number of binuclear cells.

### Statistical analysis

Statistical analysis was performed with the Student’s t-test tool on Microsoft Office Excel (version 2401). Statistical significance was assessed at a p-value < 0.05.

## Results

### Cell growth arrest and polyploidy/cell cycle changes at the wound edge – the early response

In control cells, the cell cycle exhibited a standard distribution, with most cells progressing smoothly through the G1 phase (diploid fraction) (Fig 3B and 3E). G2 and S phases were well-represented, indicating normal cell cycle progression. Upon injury, a notable shift occurred in cell cycle dynamics at 12 hours post-injury (hpi). Most cells exhibited arrest at the G1 phase, accompanied by a reduction in G2 and S phases, in stark contrast to the control (Fig 3B). Interestingly, the mononuclear polyploid cell (>4C) fraction significantly decreased from 3.88% to 0.95% (Fig 3E and 3F), while the proportion of binuclear cells tripled from 1% to 3%, with a higher accumulation at the wound edges (about 5%) (Fig 3D). This indicated the activation of a polyploidy circuit in response to the wound. Concomitant with considerable G1 phase arrest, the proportion of mitotic cells decreased twofold compared to control (from 1.45% to 0.65%) (Fig 3A) at the wound edge. At the same time, there was a fourfold increase in the anaphase+telophase/metaphase ratio (from 1.3 to 5.2) (Fig 3C), highlighting a substantial acceleration of divisions in some small proportion of cells. This early response within the first 12 hours post-injury suggests a complex interplay of cell cycle regulation in reaction to tissue damage.

**Fig 3.**
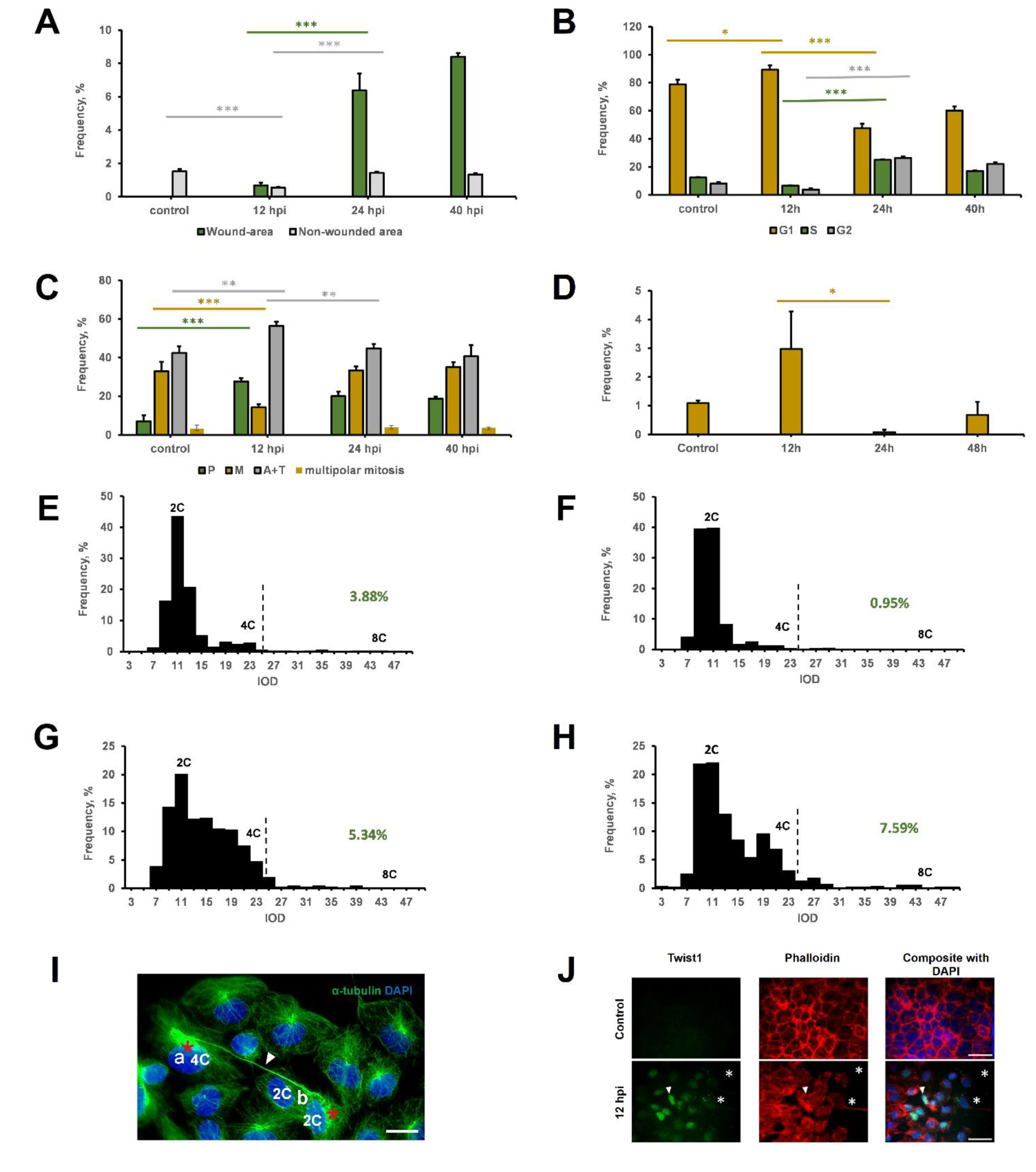
Post-wounding effects on nuclear morphology quantitative data (A-D), DNA content profiles (E-H) and immunofluorescence characteristics of α-tubulin (I) and Twist1 (J) of WB-F344 cells. (A) Mitotic indexes at different time points after wounding were separately counted within scratched and non-scratched areas. (B) Distributions of mitosis stages of cells at the edge of the wound at different time points after wounding. (C) Distributions of cell-cycle phases (G1, S, and G2) of cells at the edge of the wound at different time points after wounding. (D) Fraction of binucleated cells at the edge of the wound at different time points after wounding. Data presented as a mean ± SE from three independent experiments. ****: p ≤ 0.0001, ***: p ≤ 0.001; **: p ≤ 0.01, *: p ≤ 0.05, ns: p > 0.05. Representative cell-cycle phase distributions by in situ DNA cytometry in control (E) and within the wound edge/gap at 12 h (F), 24 h (G) and 40 h (H) after wounding. The integral optical density (IOD) values shown in arbitrary units correspond to the positioning of DNA content per cell nucleus. The dashed lines represent borders separating cell nuclei exceeding 4C DNA ploidy whose percentages are shown in green. (I) Cells at the wound edge stained for α-tubulin (green) and DNA by DAPI (blue) display two tetraploid daughter cells (“a” and “b”) whose centrosomes (shown by black asterisks) are still joined by a long bridge (shown by white solid arrowhead), which is indicative of the relatively recent cytokinesis that took place a few hours ago; however, the daughter cell “b” has already managed to perform an apparent karyokinesis, thus forming a binuclear (2C+2C) cell, whereas the daughter cell “a” remains quiescent. (J) There is no Twist1 expression in control (intact cells) with a typical “cobblestone” epithelial pattern of actin staining (red), whereas Twist1 (green) is highly expressed in cell nuclei within the wound edge especially at 12 h after wounding. Here the wound gap is shown by asterisks, and at the wound edge some cells with reorganized actin cytoskeleton have extending lamellipodia (shown by arrowhead). Scale bars = 10 µm (I) and 30 µm (J).

### Interplay of the opposite senescence and stemness regulators at the early priming phase

In the absence of injury, control cells displayed a low-level presence of the senescence marker and Cdk4/6 inhibitor p16^Ink4a^ in cell-nuclei and cytoplasm, suggesting some intrinsic balanced activity (Fig 4A). Concomitantly, the stemness regulator Nanog exhibited a moderate nuclear presence in control cells (Fig 4A). This baseline expression of Nanog in the absence of injury underscores its role as a master regulator of stemness in oval cells even under normal conditions. Upon initiation of the early priming phase following injury, both p16^Ink4a^ and Nanog displayed increased presence in cell nuclei, as evidenced by comparative immunofluorescence images. Moreover, while in control of confluent cultures a few Nanog granules were encountered in the cytoplasm, their abundance appeared in the cytoplasm and interspersed among cells at the wounded edge at 10-12h (Fig 4B). Interestingly, cells rich in cytoplasmic Nanog granules exhibited weaker nuclear staining for Nanog but stronger staining for p16^Ink4a^ compared to cells with fewer cytoplasmic granules (Fig 4B, arrowed). This observation hints at the dynamic interplay between these regulators and the potential influence of their subcellular localization on the cell’s fate. Another stemness and stress-related transcription factor and an immediate early gene response in liver regeneration, Myc was also of interest. While mostly background cytoplasm Myc- staining was seen in control WB-F344 cells (Fig 4E), at 12 hpi, at the wound edge cells, this was particularly concentrated at the centrosome area and outlined the cytoskeleton (Fig 4F), likely being presented by MycN. Its overexpression is known as arresting the centrosome cycle after DNA damage, in cooperation with wild type p53 [17]. However, the absence of Myc in cell nuclei testified to its suppressed activity as a transcription factor, in line with the decreased nuclear location of Nanog when overcompeted by cytoplasmic-nuclear p16^Ink4a^ influx. At the same time, this resulted in the Nanog protein displacement into cytoplasm as granules and conditioning of the microenvironment at a wounded edge with the stemness-self-renewal factor, which could paracrinally and autocrinally boost the next, proliferating phase of wound healing.

**Fig 4.**
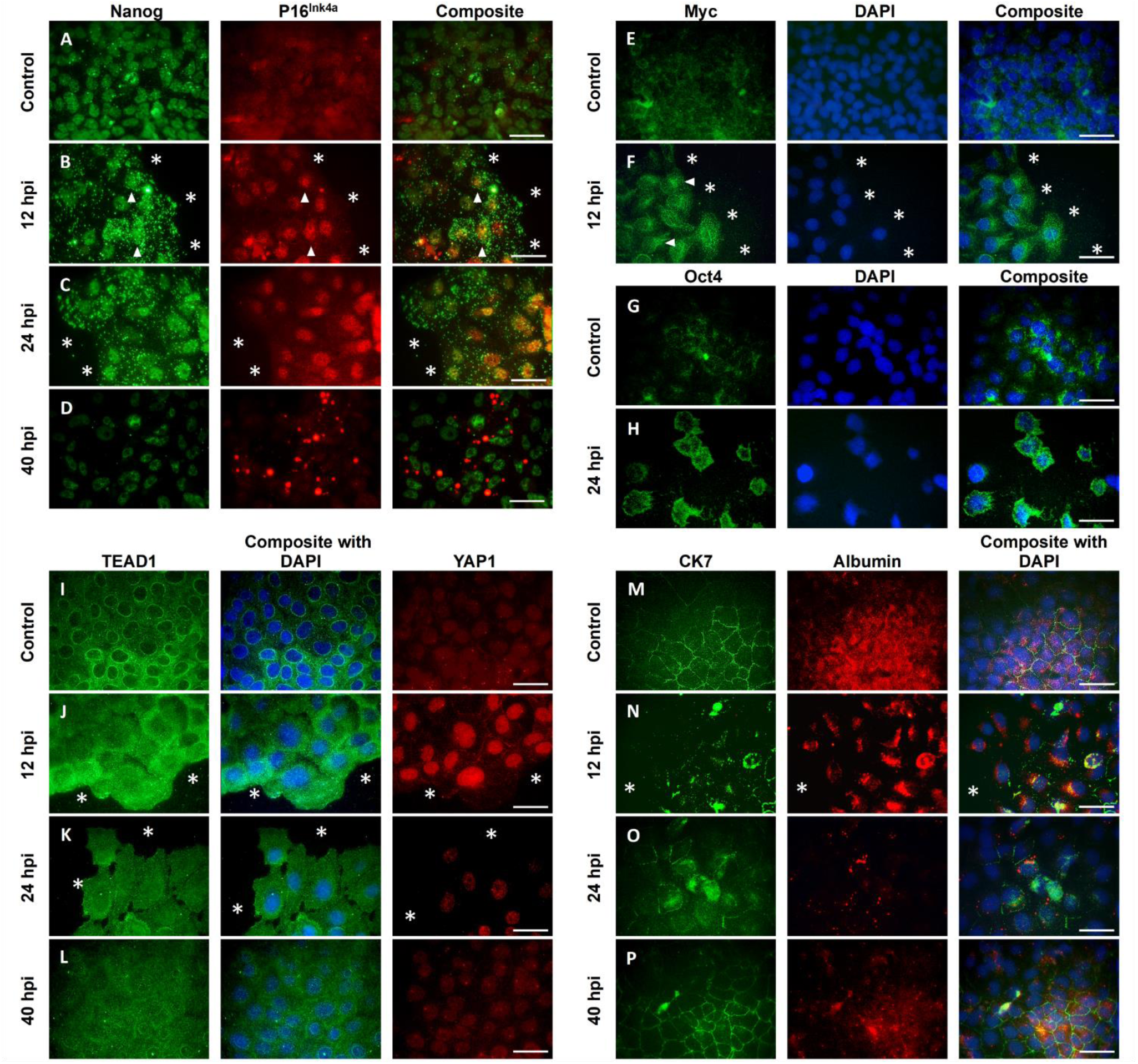
Post-wounding effects on localization, expression, and dimensional characteristics of immunofluorescence of Nanog and p16^Ink4a^ (A-D), Myc counterstained with DAPI (E-F), TEAD1 and YAP1 counterstained with DAPI (I-L), and CK7 and albumin counterstained with DAPI (M-P). Notably, at 12 h after wounding, there is an increase of Nanog cytoplasmic granules (size and numbers) accompanied by an increase of p16^Ink14a^ nuclear expression, as the white arrowheads show some most representative cases (B). In contrast, at 40 h after wounding, many variously sized 16^Ink4a^ granules drastically appear in the cytoplasm with the disappearance of Nanog cytoplasmic granules (D). At the same time, Myc at 12 h after wounding (F), compared with the control (E) increases in the cytoplasm, especially in the centrosome area (white arrowhead) of cells located closer to the wound edge. Also, at the wound edge, cells have an increased cytoplasmic Oct4 (H), compared to the control cells that express Oct4 in cytoplasm as well as in nuclei to an insignificant extent representing background levels (G). Differently localized YAP1 and TEAD1 are well expressed over an active wound healing process (J and K), pointing out the fact that after wounding (especially at 12 h; J) TEAD1 becomes markedly expressed in nuclei and cytoplasm of edge cells, and these cells are also characterized by an elevated and distinct nuclear YAP1 expression. However, these peculiarities diminish upon the wound closure time (L). Contrary to cells actively involved in wound healing, control cells have well-defined nucleo-cytoplasmic TEAD1 rings and dimly stained nuclear YAP1 (I). Albumin and CK7 drastically and synchronously change throughout wound healing (N, O, and P). Although there is an apparent difference in their localization, some shared patterns of colocalization may also be present. Further explanations are in the text. Asterisks (*) indicate the wound gap. Scale bars = 30 µm.

The change in cell ploidy and fate, motivated us to look at the Hippo pathway, an important cell and tissue growth regulator, in particular its main operators – transcription regulator YAP1 and its binding target in the nucleus, transcription factor TEAD1. Under basal conditions, control cells displayed YAP1 and TEAD1 subdued intensity and were notably absent in nuclei, as depicted in Fig 4I. Interestingly, TEAD1 revealed ring-like structures at the inner perimeter of the nuclear envelope (Fig 4I), potentially signifying lamin B1 rotation associated with accelerated cell senescence, which may be an indirect sign of the lamin B1 rotation in accelerated cell senescence and of cell nucleus turbulence observed preceding YAP1 activation [18]. As the early wound response developed, a notable reconfiguration in the Hippo pathway dynamics became apparent. At 12 hpi, cells at the wound edge exhibited increased YAP1 nuclear staining, coupled with an overall increase in TEAD1 (Fig 4J). This heightened activation indicated a responsive modulation of the Hippo pathway, implicating it in the cellular and tissue growth regulation triggered by the injury. The initial wound response also induced substantial alterations in cell phenotype, characterized by the loss of epithelial markers (CK7 and albumin), disorganization of the cobblestone-like tissue structure, and reorganization of the actin cytoskeleton (Fig 4N and 4O). Additionally, the emergence of lamellipodia (Fig 3J, arrowed) indicated a shift toward a mesenchymal phenotype, suggestive of an EMT process.

Twist1, a pivotal transcription factor associated with EMT initiation, displayed intense nuclear staining within wound edge cells at 12 hpi, in stark contrast to its absence in control cells (Fig 3J). This observation provided further substantiation for the occurrence of EMT during the initial phases of the wound response.

### Proliferation burst, MET and Nanog/p16**^Ink4a^** interplay resolution as the late wound response of WB- F344 cells

At 24 hpi there starts a tremendous elevation of mitotic activity, which continues to grow at 40 hpi (Fig 3A), 4- and 5-fold increasing mitotic index, respectively, compared with control. The proportions of metaphases and anaphase+telophase phases at these time points were comparable with the control, approximately 1.1 - 1.3 (Fig 3C). In cell cycle histograms, at 24 hpi instead of one peak at 2C and another at 4C, we found a “ladder”- like histogram starting from G1, embracing a very bulk S-fraction fusing with G2 and extending up to the hyper-tetraploid fraction (Fig 3G). More conventional histogram was obtained at 40 hpi within the cells in the damaged area with peaks at 2C and 4C, however, some “ladder” remnants could be seen (Fig 3H); the polyploidy (mononuclear) level continued to increase from 24 hpi and was 1.5-fold higher in comparison with control, also octoploid and even bigger ploidy cells are already seen (Fig 3E-3H). Dropped to about 0.2% at 24 hpi (Fig 3D) the frequency of binuclear cells may indicate their attraction to the wound, followed by segregation and use in the wound healing process occurring before the burst of mitosis.

At the time of proliferation burst (24 and 40 hpi) WB-F344 cells regained epithelial phenotype (Fig 4O and 4P), no expression of Twist1 is present anymore (data not shown) and YAP1 and TEAD1 staining became less intense, suggesting Hippo pathway tumor suppressing function resuming (Fig 4K and 4L). Interestingly, we found that Oct4, one of the main stemness factors, while weakly stained with the monoclonal antibody (covering the 1st exon) in control WB-F344 cells (Fig 4G), and at 24 hpi show the non-specific antibody binding exclusively within morphologically distinct, distorted nuclei. The absence of the nuclear Oct4 staining in our model indicates the absence of the nuclear binding and hence the transactivating activity, in line with the absence of C-Myc activity in cell nuclei (Fig 4E and 4F) and may also indicate to the restricted p53-controlled stemness potential of oval cells. The interplay between two opposites for cell fate proteins, self-renewing Nanog or senescence-inducing p16^INK4A^ continue to show heterogeneous patterns (presuming oscillations) at 24 hpi at the wound edge started early after the injury (Fig 4C), however, resolution comes at 40 hpi (almost full wound closure time) when Nanog over- dominated and “casted-out” p16^Ink4a^ from cell nuclei (Fig 4D). Previously rich in Nanog granules, cellular cytoplasm is now clear from them, however, differently sized p16^Ink4a^ positive cytoplasmic granules and aggregates, approximately 1-3 per cell (if present) appeared (Fig 4D). It could be deduced, the “bistable switch” between two toggles – stemness and senescence – in WB-F344 cells occurring along with wound healing phases provides its positive outcome.

A summary of the observed cell changes during the whole wound healing process is provided in Table 2.

**Table 2.** Summary of the observed events during the *in vitro* regeneration model.

| Cellular characteristics | Marker | Localization | Time points |  |  |  |
| --- | --- | --- | --- | --- | --- | --- |
|  |  |  | Control | 12hpi | 24hpi | 40hpi |
| Stemness | Nanog | Nucleus | + | ++ | ++ | + |
|  |  | Cytoplasm | +/- few granules | ++ granules | ++ granules | +/- |
| Senescence | P16<sup>Ink4a</sup> | Nucleus | + | ++ | ++ | - |
|  |  | Cytoplasm | + diffuse | ++ | ++ | + granules |
| Bi-potential tissue-specific stemness potency | Albumin (hepatocyte) | ++ diffuse | +/- | +/- | + | Albumin (hepatocyte) |
|  | Cdk7 (cholangiocyte) | ++ | +/- | +/- | ++ |  |
| Mobility EMT | Twist1 |  | - | ++ | - | - |
|  | Phenotype |  | Epithelial phenotype | EMT | EMT | MET |
| Hippo pathway | YAP1 | Nucleus | +/- | +++ | ++ | + |
|  |  | Cytoplasm | +/- | - | - | - |
|  | TEAD1 | Nucleus | - peri-nuclear rings | + | + | + |
|  |  | Cytoplasm | + | ++ | + | + |

Outlined is the dynamic behavior of various cellular markers associated with key biological processes such as Stemness, Senescence, Bi-potential tissue-specific stemness potency, Mobility, and Hippo pathway over distinct time points post-injury (hpi) (12, 24, and 40 hours). Each row corresponds to a specific cellular characteristic, detailing the behavior of distinct markers and their localization within the cell. The markers exhibit diverse patterns of localization (nucleus, cytoplasm), intensity (+, ++, - or +/-) indicative of their presence or activity, and distinctive cellular behaviors (e.g., granules, diffuse, transition states like EMT and MET).

## Discussion

Activation of the liver oval cells that contribute to liver regeneration is commonly observed in chronic or severe (acute) liver injury [19]. Diverse paracrine and autocrine signaling pathways have been described to control the LPC response [20]. However, it is far from being clear regarding the mechanisms that can be involved *in vivo* and *in vitro* systems for successful liver regeneration, especially in case of serious pathology. There is also an important concern about the therapy by LPCs in terms of its oncological safety. In the current study, we induced the massive injury, mechanically removing ∼70% of the culture monolayer of rat liver oval cells. In general, two phases of cell response to injury (“priming” and “proliferation”) were observed, which is in accord with the findings from *in vivo* models of partial hepatectomy [5]. Moreover, cell ploidy changes during the healing process including transient binucleation were also demonstrated, confirming the findings previously reported [13]. However, for the first time we were able to reveal and monitor the regulation of wound healing by the progenitor-driven liver regeneration involving the recurrent nucleo-cytoplasmic transfer of two opposite regulators, Nanog (stemness/cell-cycle progression) and p16^Ink4a^ (senescence/cell-cycle arrest). The upregulation of Nanog at the priming stage (10-12 hp) was converted into Nanog secretion in the form of cytoplasmic and extracellular Nanog-positive granules (vesicles) being outcompeted for the nuclear binding sites by p16^Ink4a^ transferred into cell nuclei from cytoplasm. Both regulators are known to compete for the same Cdk4/6 kinase of Cyclin D controlling the G1/S cell cycle checkpoint in stem cells: Nanog as an activator, whereas p16^Ink4a^ as an inhibitor [21, 22]. Upon completion of cell proliferation at 40-48 h post-wounding, the backward process is taking place: Nanog ousts p16^Ink4a^ from cell nucleus binding sites to cytoplasm, suppressing the excited dual state of accelerated cell senescence, which was induced by the damage. Thus, this system can be adapted to severe damage by employing a bistable switch of self-organization (schematically shown in the Graphical abstract). At present, the concept of bistable switch-based regulation is well-recognized and being intensively developed by Systems Biology as an example of adaptive thermodynamics applicable to stem, differentiated and tumor cells [23–28]. In some previous studies with breast tumor cells, there were possible fluctuations of cells with accelerated senescence induced by drugs, e.g., between the Nanog- and p16-positive, including intermediate states [26, 29]. Moreover, the experiments with knockout mice showed that senescence, in particular, its p16^Ink4a^ regulator is not only an antagonist of stemness but also is indispensable (along with secreted Il-6) to promote reprogramming [30, 31]. Although usually this feature is solely ascribed to the senescence secretome, here we demonstrate that the relationship between the potentially reprogramming Nanog and senescing p16^Ink4a^ regulators is more complex with the involvement of nucleo-cytoplasmic relocation of these two opposites, resulting in the negative feedback during second phase ending the healing process. The reprogramming potential of the epithelial oval cells (WB-F344) used in this study was also displayed by EMT-MET dynamics (as part of the same bistable switch), suggesting biphasic character of also this process. The Hippo pathway, which is known to be associated with changes of cell fate and ploidy [32, 33], has been also described in the context of liver regeneration and development [6]. We found the Hippo pathway inactive in the near-confluent WB-F344 cell culture but it can be activated by turbulence (as judged by perinuclear rings of TEAD1) and cytoplasmic-nuclear transition of activated YAP1 that coincides with the priming phase and continues during replicative stress preceding and accompanying the boost of proliferation at 24 h. The proliferation-associated “staircase” form of DNA histograms starting from the early S-phase can be suggestive of the “staggering” of replication forks caused by transcription R-loops [34]. This phenomenon is ascribed to hypertranscription activity of nuclear YAP1 that functions in partnership with TEAD1 [35].

In fact, the WB-F344 line can be referred to as somatic oval cells possessing stem-like features, such as an asymmetric cell division producing the progeny of different fates (differentiation and self-renewal). Stem cells usually reside in quiescence and proliferate under stimuli [36]. However, cell-cycle regulation of adult stem cells has not been explored as much as that of embryonic stem cells (ESCs). Most distinct features of the cell-cycle of ESCs are as follows: non-functional G1/S-checkpoint, allowing rapid proliferation, low functionality of the S-checkpoint, while the G2/M-checkpoints and spindle becomes uncoupled from apoptosis [37, 38], favoring mitotic slippage into polyploidy.

In our model, DNA histograms of the near-confluent intact control showed most quiescent cells with low mitotic index (MI) and insignificant degree of polyploidy (>4C). At the same time, there were cell-cycle progression delay at 12 hpi with a halved mitotic index accompanied by disappearance of mononuclear polyploid (>4-8C) cells and increasing of diploid (G0/G1) fraction with a negligible level of S-phase cells. Notably, at 24 hpi there was a 5-fold but moderate elevation of polyploid fraction (>4C), not exceeding 8%. That means that the G1/S-checkpoint became adapted, thereby resulting in cell-cycle acceleration, although S-phase was “staggering” at stalling replication forks due to collision between enforced replication and hyper-transcription of active genes at the beginning of S-phase [39], which is suggestive of replicative stress. In this situation, the “leakage” of the adapted G2/M-checkpoint with a lowered threshold may reduce this stress, diverting a proportion of replicated cells from the mitotic cycle into polyploidy (4C-8C). Our counts show a clear reciprocal relationship between the numbers of bi-nucleated and mononucleated polyploid cells. One could suggest that those may be playing a role of the reservoir for the emergent adaptation to wounding, thereby initially saving the energy needed for new DNA synthesis. Therefore, it seems logical to hypothesize that only these types of cells (binucleated and mononuclear polyploids) acquire the plasticity-assigned features of “avant-garde” undergoing EMT, which is necessary for initial loose filling of the wound gap creating the stromal substrate for the upcoming proliferation, simultaneously boosted by the Nanog-containing secretome. Once the aforementioned priming response is completed and binucleate cell pool becomes exhausted, the energy of mitosis due to the renewed DNA synthesis is directed to the formation of the next stock of polyploid cells (the hypothetical Scheme of the ploidy response in oval cells using the binucleation/polyploidy (2C- 4C-8C) conveyor is presented in Fig 5.

**Fig 5.**
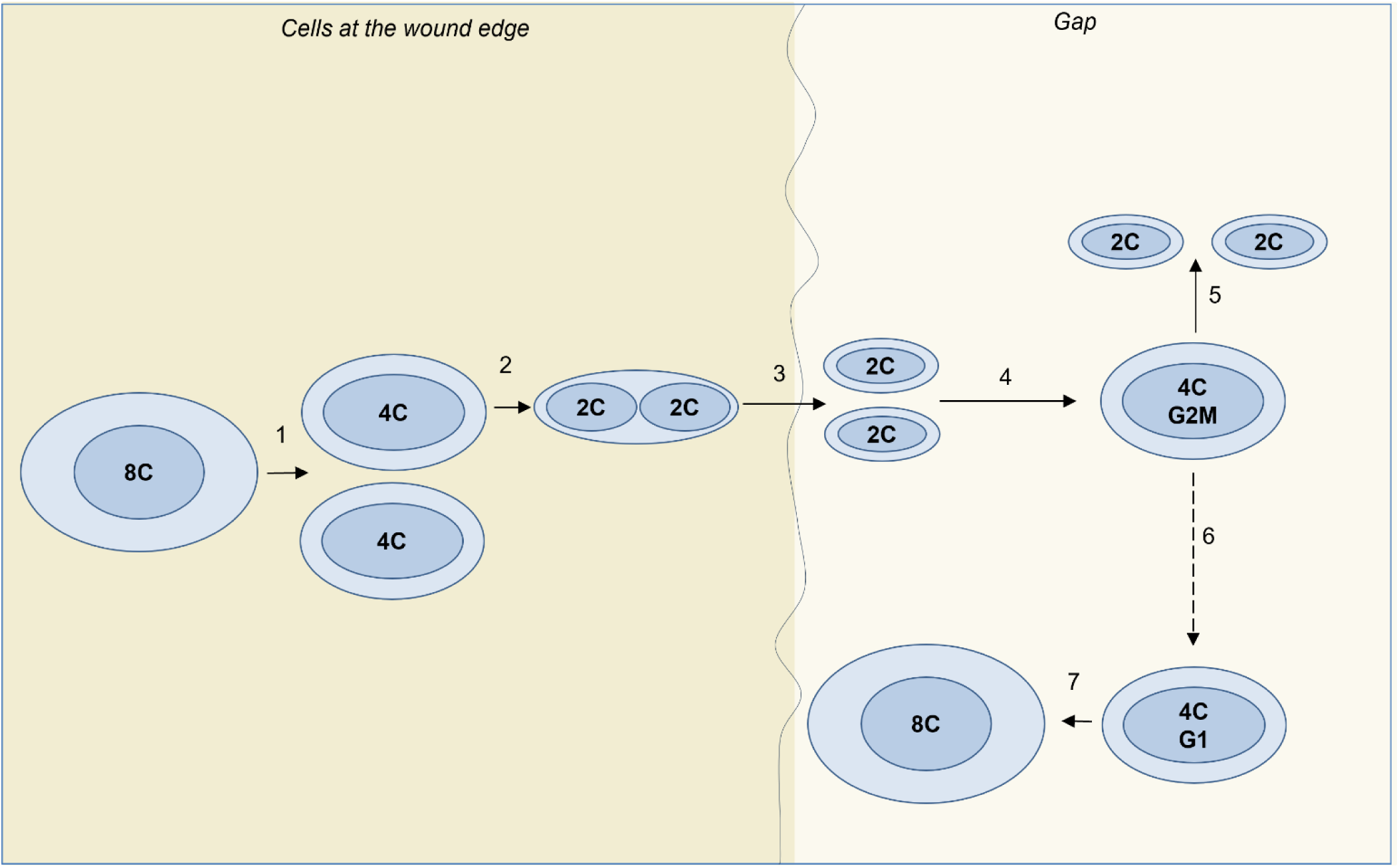
Hypothesis: Oval cells’ ploidy circuit in wound healing. (1) After the injury the fraction of polyploid 8C cells undergoes reductive divisions resulting in 4C cells, which (2) immediately perform cytokinesis- deficient karyokinesis producing binucleate cells. (3) 4C binucleate cells undergo postponed reductive cytokinesis, producing 2C daughters moving into the wound gap as mesenchymal cells via EMT. (4) In the wound gap, cells regain epithelial phenotype via MET and enter the mitotic cycle to replenish the pool of 2C cells (5). However, a portion of cells via mitotic slippage (6) advance to the new polyploidy cycle, restoring an exhausted pool of polyploid cells (7).

It is important to stress that this two-way process can be well organized by the negative feedback, thus restricting the potential of uncontrolled growth that would let the “tumor genie out of the bottle”. All toggles observed here, such as Nanog/p16, EMT/MET, and presumably binucleation/polyploidization, function here in a coordinated biphasic manner. While considering solid tumors as non-healing wounds [40], our observations, on the other hand, suggest that just wound healing by oval cells with reprogramming potential represents the barrier mechanisms against cancer. This barrier, consisting of adaptive, biphasic, and completed self-organization, is functionally efficient based on cell-cycle checkpoint regulation coupled to the nuclear-cytoplasmic shuttle of the opposite regulators. The transient activation of Myc found in centrosomes and cytoplasm but not in cell nuclei suggests that likely N-Myc with its cell migration-promoting role [41, 42] rather than nuclear C-Myc that maintains stress- dependent polyploidy linked to cancer-initiating pathways [43, 44] is functionally predominant in oval cells, thereby restricting tumor growth and uncontrolled polyploidization. Perhaps, that is why the WB- F433 cell line, relatively resistant to ionizing irradiation [15, 45], is not tumorigenic *in vivo* [14]. Further research and clinical trials are needed to use the oval cells with therapeutic purposes, offering hope for innovative and effective treatments for liver pathologies including cancer.

## Author Contributions

All authors have contributed to this article as follows:

Conceptualization, J.E. and M.L.; methodology, J.E., M.L. and K.S.; writing-original draft preparation, M.L.; writing-review and editing, J.E., B.G.; project administration, D.P.; funding acquisition, D.P.; supervision, J.E. All authors gave final approval for the manuscript.

